# Inheritance entropy quantifies epigenetic regulation of cell-cycle exit in human bone marrow stromal cells

**DOI:** 10.1101/2025.10.21.683667

**Authors:** Alessandro Allegrezza, Riccardo Beschi, Domenico Caudo, Andrea Cavagna, Alessandro Corsi, Antonio Culla, Samantha Donsante, Giuseppe Giannicola, Irene Giardina, Giorgio Gosti, Tomás S. Grigera, Stefania Melillo, Biagio Palmisano, Leonardo Parisi, Lorena Postiglione, Mara Riminucci, Francesco Saverio Rotondi

**Affiliations:** Dipartimento di Fisica, Sapienza Università di Roma, Rome, Italy; Center for Life Nano & Neuro Science, Italian Institute of Technology, Rome, Italy; Istituto Sistemi Complessi, Consiglio Nazionale delle Ricerche, UOS Sapienza, Rome, Italy; Istituto Nazionale di Fisica Nucleare, Sezione Roma 1, Rome, Italy; Dipartimento di Medicina Molecolare, Sapienza Università di Roma, Rome, Italy; Tettamanti Center, Fondazione IRCCS San Gerardo dei Tintori, Monza, Italy; Dipartimento di Scienze Anatomiche, Istologiche, Medico Legali e dell’Apparato Locomotore, Sapienza Università di Roma, Rome, Italy; Istituto di Scienze del Patrimonio Culturale, Consiglio Nazionale delle Ricerche, Montelibretti, Italy; Instituto de Física de Líquidos y Sistemas Biológicos, CONICET and Universidad Nacional de La Plata, La Plata, Argentina; CCT CONICET La Plata, Consejo Nacional de Investigaciones Científicas y Técnicas, Argentina; Departamento de Física, Facultad de Ciencias Exactas, Universidad Nacional de La Plata, Argentina; Dipartimento di Ingegneria Chimica, dei Materiali e della Produzione Industriale, Università degli Studi di Napoli Federico II, Napoli, Italy

## Abstract

Human bone marrow stromal cells (BMSC) include skeletal stem cells with ground-breaking therapeutic potential. However, BMSC colonies have very heterogeneous in vivo behaviour, due to their different potency; this unpredictability is the greatest hurdle to the development of skeletal regeneration therapies. Colony-level heterogeneity urges a fundamental question: how is it possible that one colony as a collective unit behaves differently from another one? If cell-to-cell variability were just an uncorrelated random process, a million cells in a transplant-bound colony would be enough to yield statistical homogeneity, hence washing out any colony-level traits. A possible answer is that the differences between two originating cells are transmitted to their progenies and collectively persist through an hereditary mechanism. But non-genetic inheritance remains an elusive notion, both at the experimental and at the theoretical level. Here, we prove that heterogeneity in the lineage topology of BMSC clonal colonies is determined by heritable traits that regulate cell-cycle exit. The cornerstone of this result is the definition of a novel entropy of the colony, which measures the hereditary ramifications in the distribution of inactive cells across different branches of the proliferation tree. We measure the entropy in 32 clonal colonies, obtained from single-cell lineage tracing experiments, and show that in the greatest majority of clones this entropy is decisively smaller than that of the corresponding non-hereditary lineage. This result indicates that hereditary epigenetic factors play a major role in determining cycle exit of bone marrow stromal cells.

## I. INTRODUCTION

Epigenetic inheritance, namely the transmission of phenotypic traits across generations without changes to DNA sequence, has gained great interest in biological research in recent decades [1]. Non-genetic transmission can occur through various molecular mechanisms, among which DNA methylation, histone modifications, and non-coding RNA regulation [1]; although the term “epigenetics” has drastically changed its meaning since it was coined, it is now used as an umbrella name for all non-genetic types of hereditary transmission [1, 2]. Epigenetic inheritance unfolds its full potential in the case of stem cells, as these are non-pathological cells with high replicative capacity, which makes them the ideal arena to study how different phenotypic traits emerge and evolve within a clonal population. Besides, differentiation and cell-fate, which are core issues of stem cell biology, strongly rely on epigenetic mechanisms [3, 4].

The role of non-genetic inheritance in stem cell replication has been therefore intensely studied [5–11]. It is believed that cells switch between molecularly and phenotypically distinct states that are passed on to the descendants, thus regulating their fate [9, 12]. Hence, one way to assess inheritance is to characterise cell states and track how they evolve along the lineage. Single-cell profiling allows to measure the whole transcriptome (single cell RNA-seq), as well as proteomes and metabolic signatures, hence characterising the multi-dimensional space of cell states (the ‘state manifold’ or ‘transcriptional landscape’ [13]); but this type of analysis is hard to combine with classic lineage tracing techniques, which are however important to track hereditary ramifications. A powerful alternative is provided by barcoding lineage tracing, which allows to acquire both types of information simultaneously by inferring cell state information and partial lineage relationships from bulk measurements [14–17]. Despite their great potential, barcoding techniques still present limitations in terms of applicability to human stem cells [18] and accuracy in lineage reconstruction [13].

Among the phenotypic traits that can be used to study the hereditary nature of clonal replication, *inactivity* — that we define as the state the cell enters when it stops dividing — emerges as a particularly relevant one. This is primarily for two reasons: First, because in the context of cellular aging, inactivity essentially means senescence, whose epigenetic origin has huge clinical impact [19–22]. Secondly, because inactivity is in general a more complex state than just senescence, a state that impinges greatly on the fate of stem cells and of their progenies. When a cell exits the replication cycle and stops dividing, it enters what is called the G_0_ phase [23–26]: a G_0_ cell can either be reversibly inactive (quiescent) or irreversibly so (differentiated or senescent). Hence, inactivity may be used as a proxy of that elusive holy grail of stem cell research that is differentiation potential [27, 28]. This issue is particularly relevant for bone marrow stromal cells (BMSC, also known as mesenchymal stem cells): despite being a promising source of skeletal stem cells [29], the high degree of heterogeneity of BMSC colonies severely hinders the development of clinical protocols for bone regeneration [30]. If hereditary epigenetic factors play a major role in determining activity and cell-cycle exit, they may be involved also in the regulation of BMSCs potency. Controlling the hereditary character of inactivity could therefore help developing protocols to control inter-colony heterogeneity, thus advancing the clinical potential of skeletal stem cells.

However, inactivity as a phenotypic trait has some peculiarities that make its hereditary character – if any – harder to pin down. Normally, inheritance implies that when an epigenetic change, or ‘mutation’, emerges in a cell giving rise to a new phenotype, this mutation is inherited by its progeny, thus amplifying the detectability of that phenotype with increasing generations: when enough doublings have been reached, there is sufficient bulk information in the culture to map the hereditary ramifications of that mutation through barcoding lineage tracing methods. The tricky thing about inactivity, though, is that its very impact on the colony development, namely the deletion of entire branches of the tree, is also what makes it hard to characterise it at an hereditary level through bulk methods: once inactivity emerges in one cell, this phenotype is *not* amplified in its progeny, as the effect of inactivity is rather to delete the descendants of that one cell, thus erasing all the information they potentially carried. Bluntly put, it is impossible to extract information from cells that do not exist.

Since all downstream branches of an inactive cell are missing, it seems instead that one might try to extract information from its *upstream* branches — or progenitors — which have been alive at some point in the clone development. This strategy, though, can be pursued only if inactivity emerges a few generations *after* the epigenetic mutation causing it, otherwise neither the future nor the past generations would contain any information about the emergence of inactivity. At least in the case of senescence, there is growing evidence that this is indeed the case: senescence is characterized by markers that accumulate in a gradual manner *prior* to its induction [19]; hence, it is possible that this happens more generally for the emergence of inactivity. And yet, the very absence — by definition — of the inactivity trait in the progenitors of the inactive cell, makes it unclear how to phenotypically retrace upstream the origin of the epigenetic mutation and its hereditary unfolding along the lineage. This is why current methods to search for markers of senescence resort to mass-culture molecular screening [19–22]; this type of analysis, though, is bound to miss the single-cell lineage information that is necessary to pin down directly the possible hereditary relations.

Finding a new method to back-track the ramifications of inactivity, hence establishing beyond doubt its epigenetic hereditary character at the single-cell level, is what we do here. The central idea behind this result is to extract information about the hereditary distribution of inactive cells through the calculation of the Shannon entropy associated to the topology of the entire colony. We test this method through novel experiments of high-resolution lineage-tracing of BMSC clonal colonies and find that indeed in the greatest majority of lineages the entropy is decisively smaller than that of the corresponding non-hereditary tree. Moreover, through our method, the epigenetic change giving rise to inactivity can be traced upstream, thus revealing a map of the mutation events across the colony development. This will allow us to measure the mutation lag in the hereditary structure.

## II. LINEAGE TOPOLOGY

### A. Experimental data

Human BMSCs were seeded at very low density, in order to produce clonal colonies derived only from single, spatially isolated cells; our dataset consists of 32 such single-cell-derived clones. The isolated nature of each colony is very important: if different colonies merge, their clonal nature is destroyed, severely confounding the data analysis [31, 32]. Growth was studied with phase contrast high-resolution time-lapse microscopy. Instead of adopting a fixed duration of the experiments, we fix the maximum generation of the colonies, keeping each experiment going until all replicating cells have arrived to the seventh generation, *k* = 7; in this way we do not need to correct for statistical bias [33, 34]. Details of the experiment and a full characterisation of the biological parameters of all colonies, can be found in [35].

Samples of the lineage trees are reported in Fig. 1. If no branches are interrupted, the number of cells at the seventh generation is 2^7^ = 128, and a handful of colonies do indeed reach this state (3 out of 32). But often branches *are* interrupted, which may happen for two reasons: either the cell commits apoptosis and dies, or the cell stops dividing, yet remaining alive (black capped segments in Fig.1); in this last case the cells enters the G_0_ phase. We classify a cell as G_0_ if it does not divide for 84 hours (3.5 days), at which point we stop tracking it; this threshold is to be compared to the mean division time of BMSCs, which is 20 ± 4 hours (the robustness of this G_0_ criterion is thoroughly tested in [35]). Only 17 cells die in our whole dataset, against 310 G_0_ cells; given such paucity of apoptosis and given that the effect of a dead cell on the lineage topology is exactly the same as a G_0_ cell, we will call *inactive* the whole set of G_0_ and dead cells.^1^ Conversely, we will call *active* those cells for which we do observe a mitosis. The final ‘leaves’ of the tree, i.e. cells that belong to the last generation (*k* = 7), are only tracked up to their birth, not up to their division, hence they are neither labeled as active nor inactive; we call these cells the *k*_7_ leaves of the lineage tree.

**FIG. 1.**
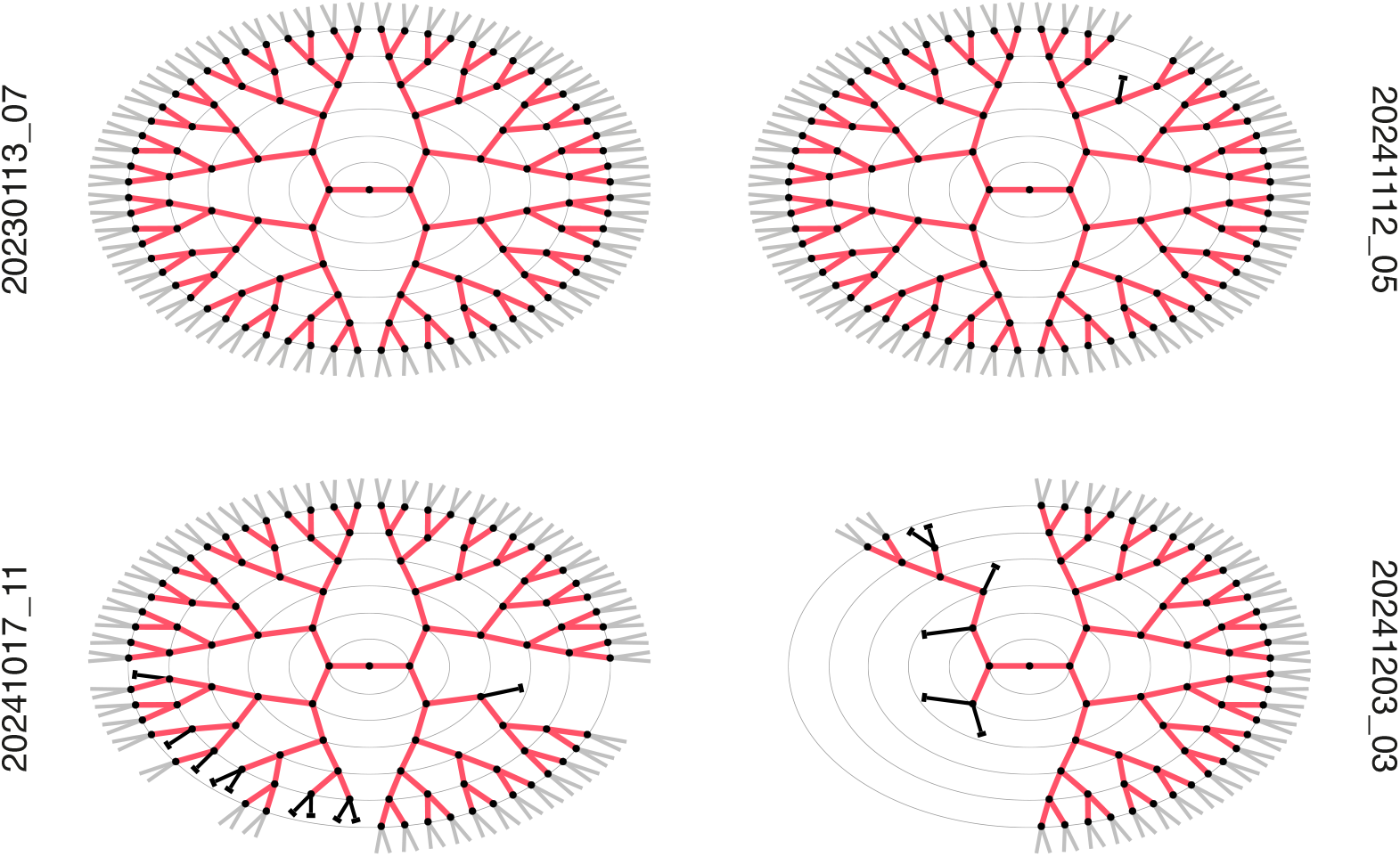
Lineage trees of four samples of BMSC clonal colonies. Red segments represent active cells and black discs represent mitosis; the length of each segment is fixed, not related to the division time. Inactive cells (which are all G_0_ in these samples) are represented as black segments ending in a cap. The first cell of the colony is not represented, as we do not record its birth, but only its division (the central disc of the tree). Ovals separate different generations. Cells at the last generation (the *k*_7_ leaves), are tracked up to their birth, not up to their division, hence they are neither labeled as active nor inactive and we represent them in grey.

### B. The peculiar distribution of inactive cells

Inactive cells determine the topology of a lineage through both their number and their position within the tree. The position of an inactive cell can be described by two ‘coordinates’: its generation *k* and its radial position. The first impacts greatly on the number of missing *k*_7_ leaves: the earlier the generation at which a cell becomes inactive, the larger the number of leaves that are cut; for example, the effect of a G_0_ cell is very different if it is located at generation *k* = 2 (32 missing *k*_7_ leaves) or at *k* = 6 (2 missing *k*_7_ leaves). At first sight, this effect seems related to the Luria-Delbrück argument about the uneven impact of hereditary mutations [39]: if the probability of a mutation is homogeneous throughout the lineage, the fluctuations in the occurrence at different generations causes very large heterogeneities in the final number of cells carrying the mutation, which is the conceptual basis for all Luria-Delbrück-inspired fluctuation assays [40]. However, because in this case the phenotype is “inactivity” and because the very definition of inactive cell is that it does not have any progeny, the fact that the inactive phenotype is inherited by the (non-existent) descendants of the inactive cell is completely trivial and thus uninformative. Therefore, to check whether or not the mechanism that ultimately leads to inactivity has an hereditary nature, we cannot use a standard fluctuation assay. Instead, we have to employ the lineage information to check whether some kind of mutation occurred *before* the emergence of the inactive cell, namely upstream in the tree.

To make progress, we notice that trees with many inactive cells display something more than the mere amplification of fluctuations due to the different intergeneration placement, namely the fact that the radial distribution of inactive cells across different branches has quite a nontrivial pattern. Consider for example lineage `20241017_11` in Fig. 1: there is clearly something peculiar about the locations of the inactive cells in this tree, as they seem more frequent within the same few branches on the south-west side in a definitely non-random way; a similar observation could be made for lineage `20241203_03`, where inactive cells are mostly concentrated in the west wing. This is a very general trait in our dataset: in most lineages we observe significant non-random heterogeneities in the number of inactive cells between branches, indicating that the emergence of these cells is more likely in some branches than in others. If we follow a branch with many inactive cells upstream along the lineage, an hereditary mechanism would suggest that at some point we must find a mitosis at which some factor linked to the probability of inactivity changed, making one of the two daughters more prone to producing inactive cells in its progeny than the other. We need to put this hypothesis to a quantitative test, and to do that we will use the concept of *entropy*.

## III. INHERITANCE ENTROPY

### A. Inactivity imbalance

As we have noted, the most conspicuous effect of the presence of an inactive cell is that all cells that would have been born as its progeny are instead missing, so that the number of cells at the final generation, *k* = 7, is smaller than what it could have been. Hence, we will use the number of missing *k*_7_ leaves as an indicator of the impact of inactive cells within that branch. We proceed as follows.

Each mitosis in the lineage generates two branches, left and right; in a mitosis connecting generation (*k* − 1) to generation *k*, each branch can potentially produce up to 2^(7−*k*)^ descendants at generation 7. However, if inactive cells emerge downstream along these branches, the *actual* number of *k*_7_ descendants will be smaller (some of these leaves will be missing). Given a mitosis *m*, we will indicate with 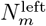 the number of missing *k*_7_ leaves in its left branch and similarly for 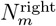 (see Fig. 2). We stress that the count of missing cells is done only among the cells of the last recorded generation (in our experiments, the *k*_7_ leaves).

**FIG. 2.**
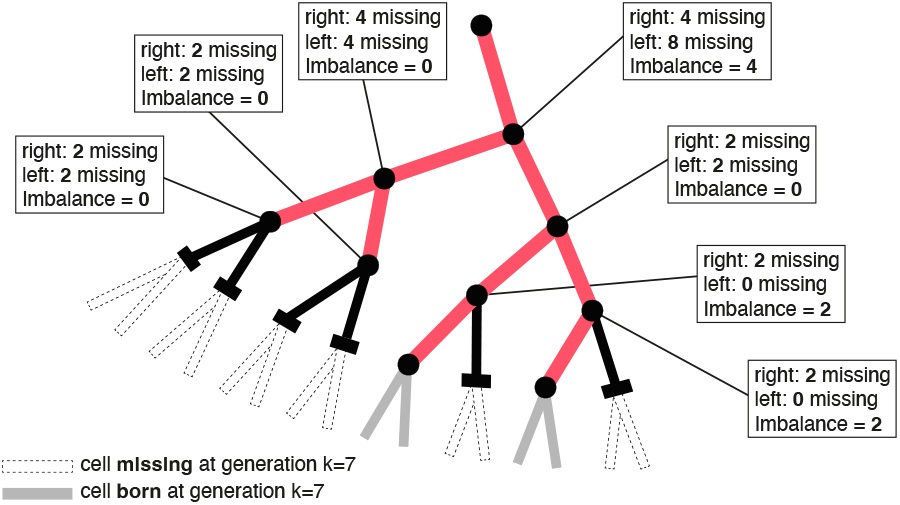
Illustration of how the inactivity imbalance is calculated, here using a portion of a real lineage (`20230503_13`). White dotted segments represent *k*_7_ leaves that are missing because of the presence of some inactive cells upstream in that branch. For each mitosis/node, the inactivity imbalance is defined as the modulus of the difference between the number of missing *k*_7_ leaves in the right branch and the number of missing *k*_7_ leaves in the left branch.

We then define the *inactivity imbalance* of mitosis *m* as the difference between the number of missing *k*_7_ cells within its left and right branches (see Fig.2),

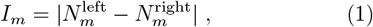

where we use the absolute value because the actual left-right labelling of the branches after a division is arbitrary.

The interesting thing about the inactivity imbalance is that it provides a sort of ‘mutation’ map, giving a specific information about what happens at each mitosis along the tree (see Fig. 3). A large imbalance on a certain node means that the two branches generated by that mitosis are *ultimately* very different from each other, suggesting that a ‘mutation’ in the probability of generating inactive cells occurred there. At the level of the analysis performed here, we cannot know whether this mutation occurred *at* the mitosis itself, or in one of the two sister cells *after* it was born out of that mitosis, as these two cases are indistinguishable from a topological point of view; just as a matter of convention, in the following we will attribute the mutation to the mitosis. The crucial point is that the inactivity imbalance provides the information about a possible change in the propensity to inactivity *before* — namely upstream in the tree — the inactivity phenotype is actually expressed. The imbalance is therefore a promising observable in the search of an hereditary test of cell-cycle exit.

**FIG. 3.**
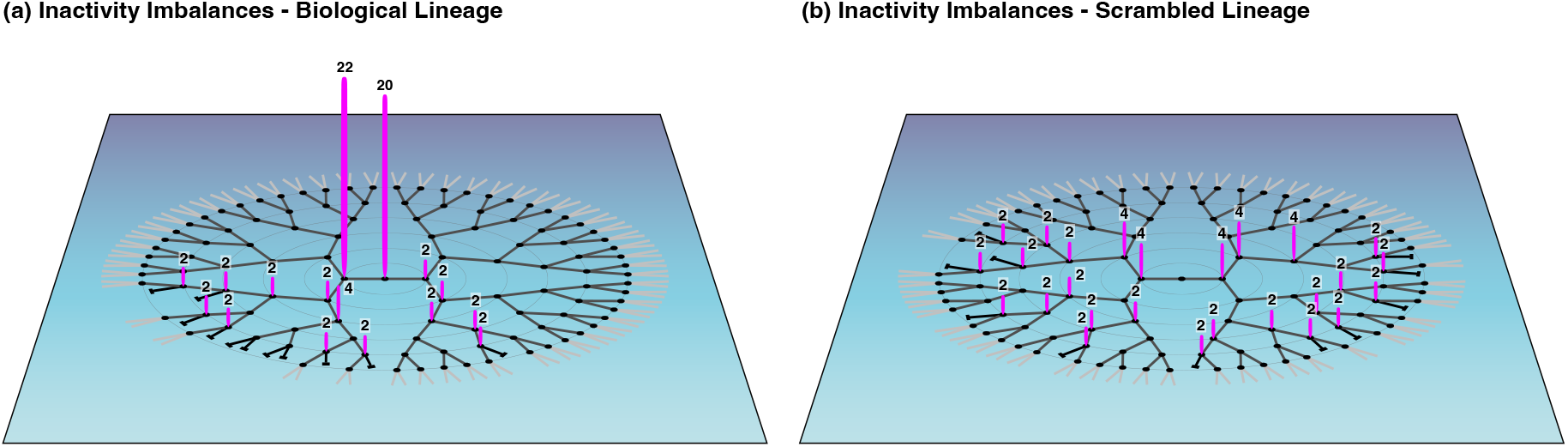
The distribution of the inactivity imbalance across the different mitosis is very different in biological vs. randomly scrambled trees. **(a)** For lineage `20230503_13`, the inactivity imbalance, *I*_*m*_, is graphically represented as a purple vertical brushstroke on each mitosis/node (null imbalances are not marked). In this lineage most of the imbalance is concentrated on just two nodes; of these two mitosis, the one connecting generation 1 to generation 2 (imbalance 22) is most likely that at which there has been a mutation in the probability of emergence of inactive cells. This large heterogeneity in the distribution of the imbalances across the lineage is the clearest symptom of inheritance. The entropy *S* measures exactly this heterogeneity; in this lineage *S*_biological_ = 2.18, which is rather low compared to: **(b)** One instance out of the 10^6^ randomly scrambled versions of lineage `20230503_13`; the number of inactive cells and their generation is the same as in the biological lineage, but their radial positions across the tree have been randomly reshuffled, hence severing all potential hereditary relationships. In this tree the imbalance is more homogeneously distributed across the nodes than in the biological tree, thus giving a much lower inheritance signal: the entropy is indeed significantly larger than the biological one, *S*_scrambled_ = 3.17. If we repeat the scrambling 10^6^ times, in only 7 cases we find *S*_scrambled_ ≤ *S*_biological_, proving that the evidence of inheritance is very strong in this clone.

### B. Entropy

At variance with other measures of tree imbalance, such as the Colless index [41, 42], we do not consolidate the inactivity imbalance (1) into a sum over all nodes of the tree. This is a crucial point: when trying to characterise inheritance, a large inactivity imbalance at a few nodes is *not* the same as a buildup of small inactivity imbalances scattered over many nodes; if there were no inheritance, but just a uniform non-hereditary probability of emergence of inactive cells, the tree would anyway show small random imbalances scattered all over the lineage. Instead, inheritance is signalled by an exceptional event, namely a large imbalance concentrated on one or a few nodes, revealing a difference between the two cells generated at that mitosis, which is passed on to their progenies (see Fig. 3). Therefore, we need a quantity able to reveal when there are atypically large values of the inactivity imbalance in the lineage, and when — on the other hand — the imbalances are all more or less the same, scattered across the tree. The most promising tool to do this is the Shannon entropy. We first define the *normalized* inactivity imbalance of mitosis *m*,

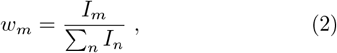

where the sum is extended over all mitosis in the lineage. Then, using the weights *w*_*m*_, we define the colony inheritance entropy as,

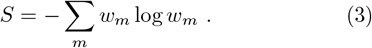

The entropy *S* is high when the inactivity imbalances *I*_*m*_ all have approximately the same value, forming a random pattern across the tree, so that we receive no particular information about the occurrence of an inherited change at any mitosis in the lineage; on the other hand, entropy is low when there are few large imbalances concentrated on a handful of mitosis, as in that case we have reliable information that an inherited change occurred at those mitosis (Fig. 3). In short — and quite coarsely — high entropy means inheritance is unlikely, while low entropy means strong evidence of inheritance. As we shall see in the next Section, though,”high” and “low” entropy will only be significant in a comparative way.

We stress that the entropy in (3) is *not* the entropy associated to the probability distribution of the inactivity imbalance, because the weight *w*_*m*_ is not the probability of a certain value of *I*_*m*_, but the (normalized) imbalance itself; indeed, the sums do not run over all possible values of the imbalance, but over all possible nodes (i.e. mitosis) where the imbalance is defined. For the same reason, the normalised weights *w*_*m*_ in (2) do not have an obvious frequentist interpretation; therefore, we should not blindly rely on the standard properties of *S* in the context of Shannon information theory. It is also worth noting that similar versions of the entropy have been employed in at least two different contexts: in astronomy an analogous quantity has been defined from the intensity field of galaxy radioastronomical data for image reconstruction [43]; in economics and social sciences, the entropy in (3) is related to Theil’s index of economic inequality [44]. In all these contexts — including ours — the key idea is to pinpoint the anomalous values of some observable (inactivity imbalance, signal intensity, economic imbalance) across a certain “space” (lineage, image, country), rather than across a certain probability distribution. This is what the entropy defined in (3) does.

### C. Inheritance test

The inheritance entropy of a lineage depends first of all on the specific arrangement of the inactive cells, but it also depends on their total number and on the overall size of the tree. Therefore, it is impossible to establish what are high or low values of *S* on an absolute scale; but fortunately we do not need to know that, because we are not attempting to compare different biological lineages using *S*. Instead, we want to know how likely it is to obtain the same entropy as a specific biological lineage, in a tree where all potential inheritance relations between mother and daughters have been severed. In other words, we want to compute the P-value of the biological entropy we measure to the null case of a completely *non-hereditary* tree.

This task is easily achieved by randomly scrambling the actual biological tree: at each generation *k* all cells are randomly reshuffled *within that same generation*, and when two cells are exchanged, so are their progenies; then we move to generation *k* + 1 and we scramble again, and so on (see Appendix A for a detailed description of this procedure). Notice that in this way potential hereditary connections are randomized, and yet the number of inactive cells at each generation remains the same; therefore, the scrambling procedure increases the entropy by erasing hereditary information and producing a completely non-hereditary tree with the same number of inactive cells per generation as the original biological tree (see Fig. 3, Fig. 4 and Fig. 7). The inheritance entropy of this scrambled tree is measured and then the scrambling procedure is repeated one million times, to produce a non-hereditary ensemble. The P-value is finally given by the fraction of scrambled trees that happen to have *S*_scrambled_ ≤ *S*_biological_, namely by the probability that a completely non-hereditary tree gives an evidence of in-heritance stronger than or equal to the biological tree. In this way, the highest significance of the test is P < 10^−6^, which happens when none of the 10^6^ reshuffled trees has an inheritance entropy smaller than the biological one.

**FIG. 4.**
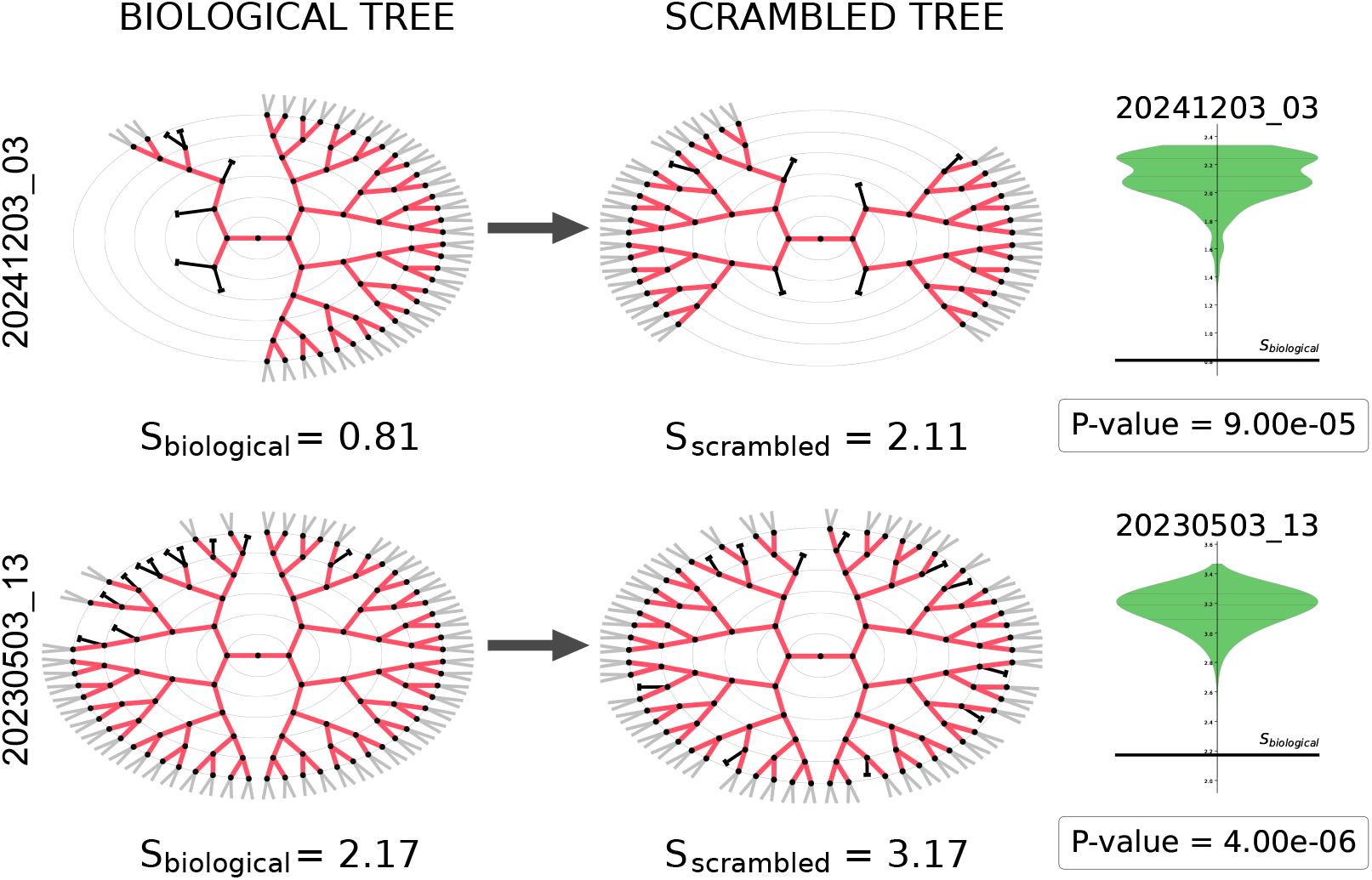
Two examples of how the hereditary structure of the inactive cells is erased by the scrambling procedure and how this impacts on the inheritance entropy. **Top:** Lineage `20241203_03` (left) has a strong concentration of inactive cells, and therefore of missing cells, in the west wing of the lineage; in this tree the imbalance is generated at the very first mitosis (the central node). Instead, a randomly scrambled version of the biological tree (center) has inactive cells distributed evenly across all branches. The entropy of the biological tree is *S*_biological_ = 0.81, while the entropy of the corresponding scrambled tree is *S*_scrambled_ = 2.11. When we produce 10^6^ scrambled versions of `20241203_03`, we very rarely find *S*_scrambled_ ≤ *S*_biological_; this means that the probability that the entropy of the biological case is so small by pure chance is extremely low. The distribution of the randomly reshuffled entropies is shown as a violin-plot on the right: it is evident that the biological value of the entropy is way below the bulk of the distribution of the reshuffled entropies. The P-value is simply the integral of this distribution below *S*_biological_, which gives P= 9.0 × 10^−5^, making the inheritance test for colony `20241203_03` highly significant. **Bottom:** A similar situation holds for lineage `20230503_13`, which also gives a very strong inheritance signal.

The outcome of the inheritance test on our BMSC colonies is reported in Fig. 5. We have 32 colonies in total in our dataset; in 4 colonies the test cannot be run because they have either zero or one inactive cell, hence the scrambling procedure produces always the same tree. Of the 28 colonies for which we can run the test, 21 (75%) have lineages with a significant hereditary character (P < 0.05), while in 7 colonies (25%) the inheritance test gives a non-significant result (P ≥ 0.05). The non-significant cases are mostly given by colonies with a very small number of inactive cells, for which the inheritance test — which is based on scrambling in many *different* ways the positions of the inactive cells — is not expected to be very effective: the four colonies with P> 0.1 have either 2 (`20241112_ 02, 20250328_16`) or 3 (`20241017_16, 20250328_02`) inactive cells; two colonies with 5 (`20241203_09`) and 7 (`20230113_08`) inactive cells have 0.05 < P < 0.1; in just one colony in our dataset (`20240627_06`) the inheritance test gives a non-significant result (P= 0.13) despite having 22 inactive cells. In conclusion, the overall result of the inheritance test indicates that the topology of the largest majority of human BMSC colonies cannot be attributed to random variations in the positions of inactive cells across different branches, thus proving that the processes responsible for regulating the probability of emergence of inactive cells have a strong hereditary character (in Appendix B we demonstrate that the structure of inactive cells cannot be due to physical proximity). Given the short timescales of the experiment and the low number of generations, it is highly unlikely that such structure is determined by genetic mutations [10, 32]; instead, our results are most likely due to the hereditary propagation of non-genetic, that is epigenetic, factors regulating cell-cycle exit.

**FIG. 5.**
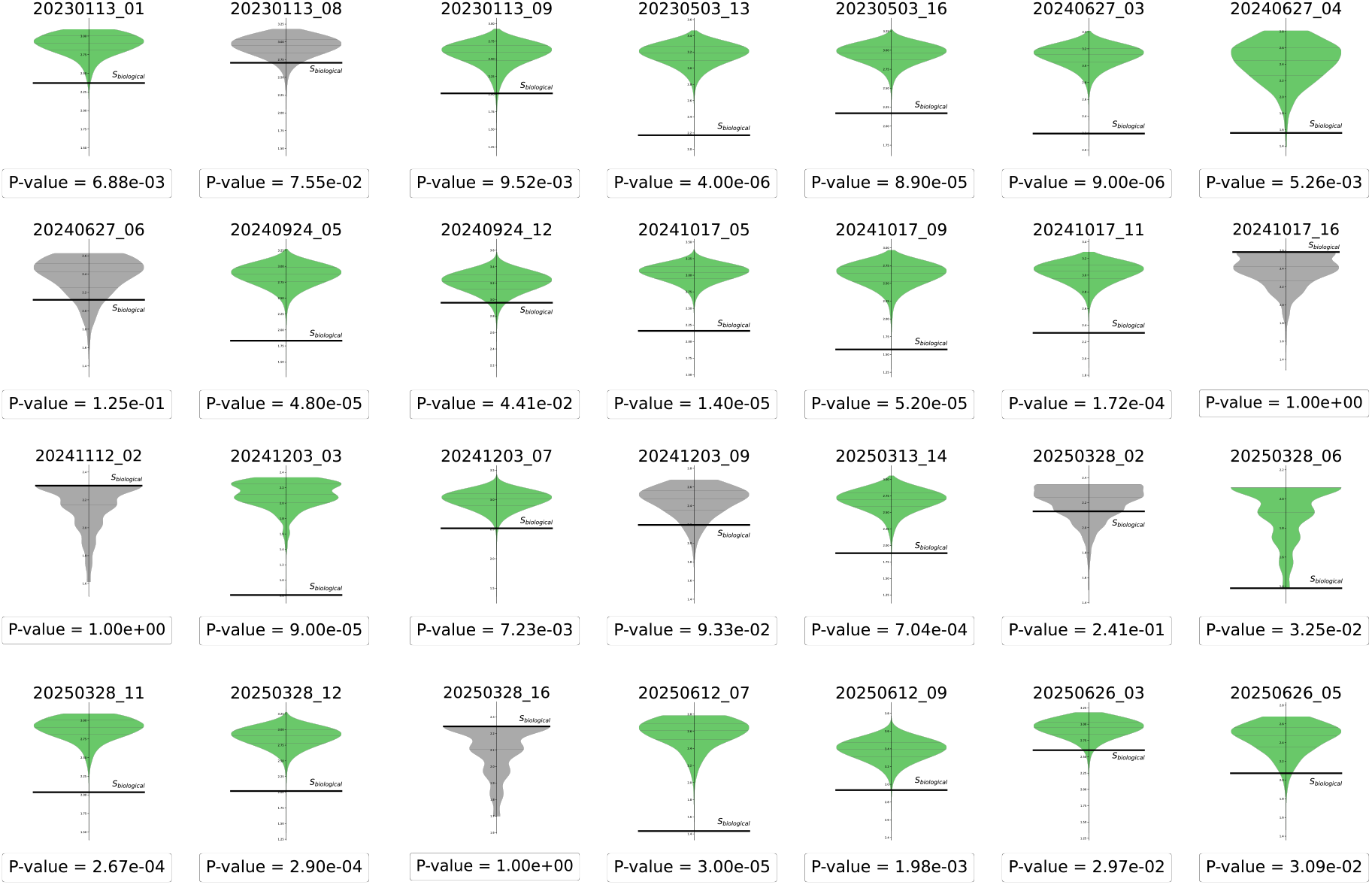
Result of the inheritance test for all colonies in the dataset for which the test can be run (i.e. for lineages with more than one inactive cell). For each colony, the violin plot represents the probability distribution of the entropy for the set of 10^6^ randomly scrambled trees, while the black line is the value of the entropy of the non-scrambled biological tree. The P-value is the total area of the violin plot which lies below the black line. Green: significant result of the inheritance test (P-value < 0.05). Gray: non-significant result of the inheritance test (P-value ≥ 0.05).

It is important to stress that a non-significant result of the test *does not* imply that there is no inheritance at work, but merely that there is no detectable change in the probability of emergence of inactive cells along that particular lineage: the lack of variation in the expression of a character makes inheritance undetectable, not necessarily absent; hence, the test is a sufficient, but not necessary, condition for the presence of an hereditary mechanism regulating inactivity. Were we to consider more complex factors, it could very well turn out that a lineage that did not pass the inheritance test at the bare topological level analysed here, still results hereditary according to these other factors. But we cannot help noticing that calculating the entropy and its P-value is easy and quick, as one needs nothing but the bare topology of the lineage. Hence, it seems lucky, and interesting, that most BMSC colonies display a sharp hereditary character already at such a basic level as the bare lineage topology, namely in relation to the quite fundamental cell state of activity vs inactivity.

### D. Navigating the imbalance map

As we have already remarked, the value of the inactivity imbalance defined on each node of the tree, i.e. on each mitosis of the colony, provides a sort of map of the most likely sites where mutation events occurred across the tree. We have to be careful about one point, though: when there is a chain, crossing different generations, of subsequent nodes all with similar anomalous imbalances, it is the node at the *latest* generation (largest value of *k*) that actually corresponds to the mutation event.

In order to understand this point let us go back to Fig. 3. In this highly hereditary lineage, the largest part of the imbalance is concentrated on just two nodes: *i)* the central mitosis, which gives rise to the two cells of generation 1, let us call this node M1; *ii)* and one of the two mitosis separating generation 1 from generation 2, let us call this node M2. We claim that the actual mutation occurred in M2, while the spike in M1 is simply a byproduct of that in M2. The reason is that the significance of a spike in the imbalance must be assessed going *upstream* in the lineage, starting from the terminal cells, i.e. the *k*_7_ leaves. If we do this, we see that the first anomalous node that we meet is indeed M2: something clearly happened at this mitosis, because its right branch (no inactive, nor missing cells) is completely different from its left branch (which instead has 10 inactive cells, causing 22 missing cells), and this large left-right asymmetry occurs for the first time at this node, when we navigate the tree upstream. On the other hand, if no other mutation occurs at M1, this node would still display roughly the same large imbalance as M2, up to small random fluctuations, which is exactly what happens in this colony.

Reversing the order of the argument, i.e. starting from the central node and navigating the lineage *downstream*, each time we find a mitosis with a very large anomalous imbalance, we can only conclude that in one mitosis of its descendants there is a mutation. It is therefore important to avoid giving to the inactivity imbalance any temporal interpretation: it is a quantity that has a value and a meaning only once the entire colony has been traced up to a certain generation. If, in the colony of Fig.3, we traced one more generation of cells, the imbalance could change on *all* nodes, even those at the early generations; this is not surprising, as in the colony traced up to *k* = 7 there might have already been some mutations that will only become visible in terms of inactive cells once we trace one or two extra generations beyond the 7^th^.

### E. The inactivity mutation lag

Once we have established a procedure to identify the mitosis where there is a change in the probability of inactivity, an inspection of the imbalance map in those lineages that pass the inheritance test shows that the mutation ultimately leading to the anomalous accumulation of inactive cells within a branch, typically happens 3–4 generations before these cells start emerging in that branch (Fig. 6). Inactive cells may either gradually appear in the ‘mutated’ branch across different generations (as in `20230113_07` or `20241203_03`), or they may show up all at the same generation (`20250328_11`); moreover, in some clones there are two independent mutation events (`20240627_03`). Due to this complex scenario, there might be more than one way to quantify this inactivity mutation lag; if we stick to the simplest definition and count how many links separate the mutation from the *first* inactive cell emerging after the mutation, we obtain that the average value of the lag over all BMSC clones passing the inheritance test is 3.1 generations. The existence of this mutation lag indicates that what we hypothesised in the Introduction, namely that the epigenetic change responsible for the emergence of inactivity occurs a few generations *before* inactivity itself is expressed, is actually true. As we have already noted, this phenomenon is consistent with results reported in the literature about cellular senescence [19]; in a different context, the presence of generation delays between cell fate decisions and the onset of lineage markers has been found in hematopoietic stem cells [45].

**FIG. 6.**
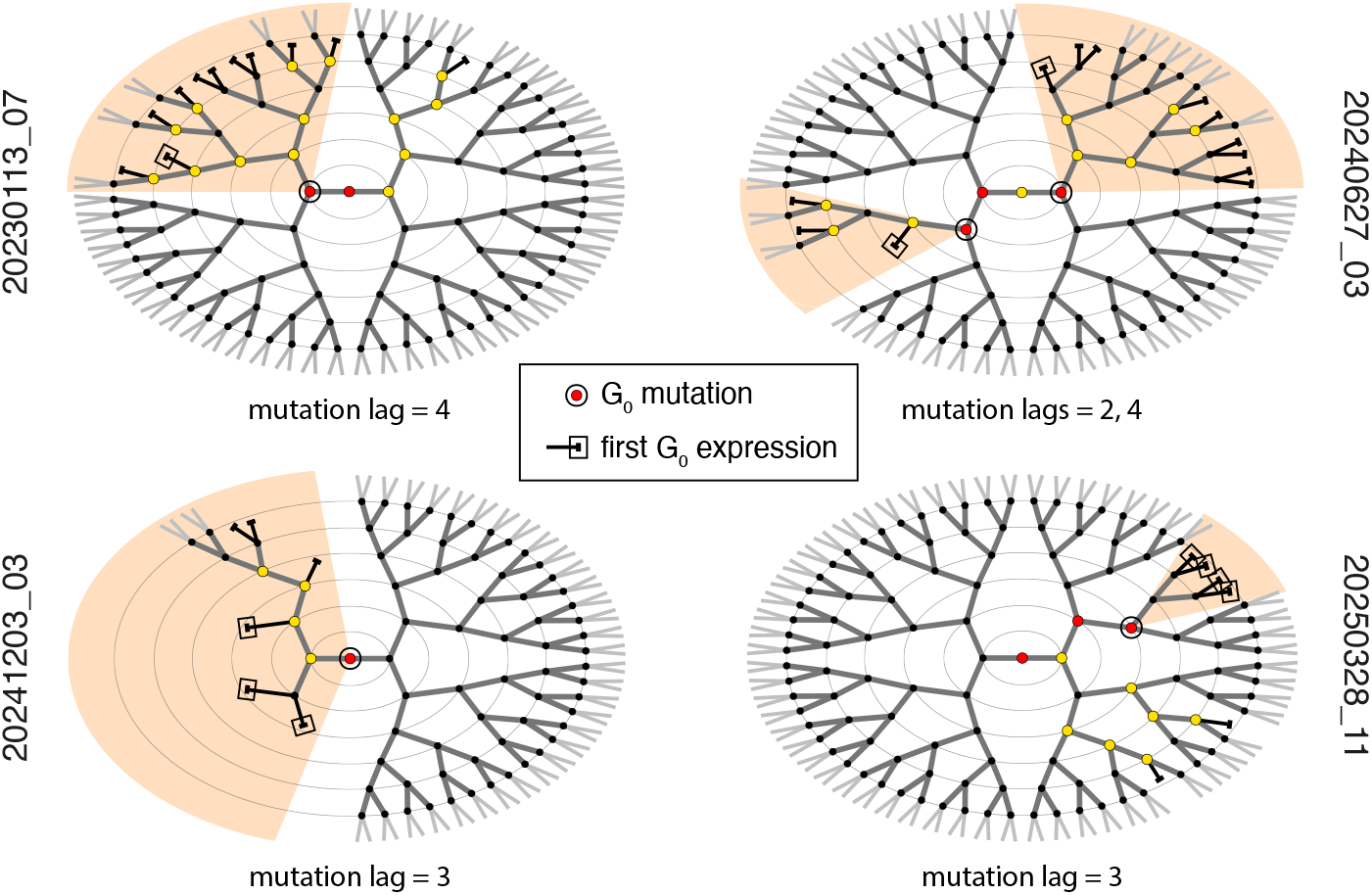
Visualisation of the inactivity mutation lag in four BMSC clones passing the inheritance test. Each mitosis *m* is coloured according to its normalized imbalance: *w*_*m*_ = 0 – black; 0 < *w*_*m*_ ≤ 0.15 – yellow; *w*_*m*_ > 0.15 – red. The mitosis whose anomalous imbalance signals a mutation in the probability of inactivity is marked by an extra external black circle and the shaded area highlights the branch impacted upon by that mutation. The *first* G_0_ cell in the mutated branch is marked by an extra external black rectangle. The mutation lag is simply given by the number of links – i.e. generations – separating the G_0_ mutation from the emergence of the first G_0_ cell. In colony `20240627_03` there are two independent mutation events, with lags 2 and 4. Colony `20250328_11` is a more complex case: it seems that there are two concatenated mutation events, one at the very first mitosis in the lineage, differentiating the left and right wings of the tree, and another (stronger) event separating generation *k* = 3 from *k* = 4 (the one highlighted in the panel); a more in depth analysis is needed to work out whether these two events are correlated.

**FIG. 7.**
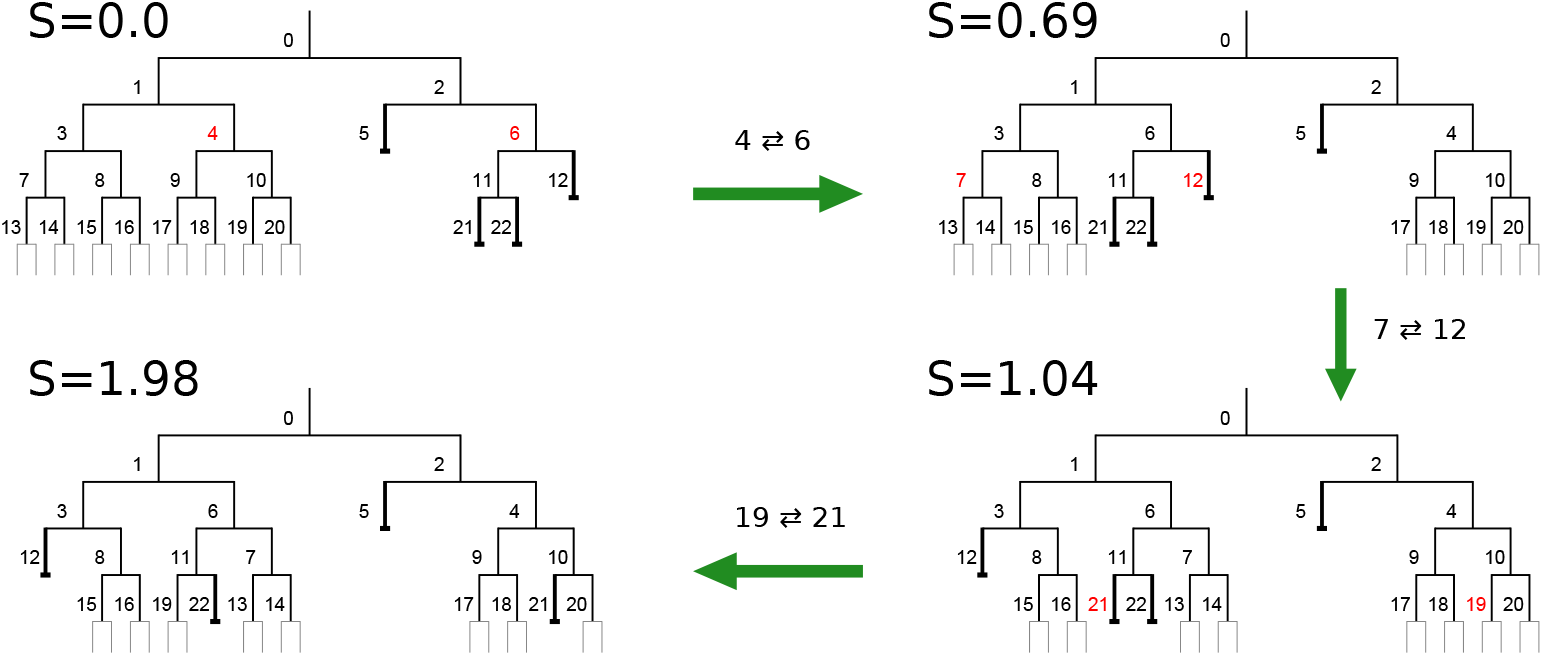
Illustration — using a fictitious tree — of the process of lineage scrambling used in the inheritance test. This tree (up-left) is designed to have a clear clustering of G_0_ cells on the right wing, and therefore a low entropy (in this case the lowest value, *S* = 0 — the reader is invited to check that there is no arrangement of inactive cells that has a more “ordered” structure than this). We select randomly two out of the fours cells at generation *k* = 2 (those with red label) and we switch them (switching the two cells at generation *k* = 1 produces the same lineage); in so doing, we carry with each cell also all its descendants. We then pass to generation *k* = 3 and randomly switch two cells and finally do the same at generation *k* = 4. With each step of scrambling the entropy grows, and in the end a tree is obtained with a much higher entropy than the original one (*S* = 1.98), which is why we can conclude that the inheritance signal of the starting lineage is very significant.

The existence of a lag between the epigenetic change leading to inactivity and its manifestation, is reminiscent of the *incubation period* introduced by the commitment theory of cellular aging [36–38], whose goal was to model the evolution of senescence in human diploid fibroblasts. There are differences with our findings, though: according to [36, 37], after the incubation period, *all* the descendants generated by the committed cell die out, while this is clearly not the case in our data. Instead, BMSC lineages suggest that at the mutation there is no deterministic commitment to inactivity, but rather a sudden increase in the probability to produce inactive cells; this probability is passed-on in a hereditary way and it acquires a phenotypic impact only a few generations later, when enough cells have been generated to produce some inactive descendants. We notice that our method is able to capture the progression to inactivity (and quite possibly senescence) on the short time scales of the first proliferation stages of the clonal colony, contrary to most studies, which rely on mass population analysis across multiple passages and focus on the long time scales [46].

Finally, we stress once again that it is only thanks to the inactivity mutation lag that the hereditary nature of cell-cycle exit could be established: if the inactive cell was born *exactly* at the same mitosis where the epigenetic change responsible for inactivity occurred, it would be impossible to assess inheritance. This seems at odds with the fact that – lag or not – if the number of *k*_7_ leaves cut by an inactive cell is large, the imbalance will also be large, thus lowering the entropy, which should give a strong inheritance signal. In fact, the situation is more subtle. Imagine a tree in which there is just one mutation and imagine that that mutation occurs at the central node of the tree, the one producing the two *k* = 1 cells; imagine also that *there is no mutation lag*, hence (say) the left cell becomes immediately inactive, thus cutting all its 64 potential *k*_7_ leaves, while its sister – the right cell – generates a full half-tree, ultimately producing all its 64 *k*_7_ leaves. In this case, a large inactivity imbalance, *I* = 64, is entirely concentrated on just that central mitosis, which gives a very small entropy, in fact the smallest, *S* = 0. However, when we attempt to randomly reshuffle this tree, we immediately recognise that the only move we can make is to switch the two *k* = 1 sisters, which produces exactly the same lineage; hence, *all* random trees have *S* = 0, so that the P-value is equal to 1 and we have zero inheritance signal. This is a general phenomenon: the absence of a mutation lag would imply that the locations of inactive cells coincide with the locations of the mutation events; if these events are uncorrelated from each other (and there are no strong reasons to think otherwise), then reshuffling the inactive cells creates the same statistical lineage and the entropy is not lower in the random set, on average. This argument highlights very vividly that the *absolute* value of the entropy does not have any meaning; its only its value *relative* to the random set of lineages that is significant: a lineage is highly hereditary not because its entropy is low, but because it is *lower* than that of the corresponding non-hereditary ensemble. In this respect, the reshuffling procedure is as crucial an ingredient of the inheritance test as the entropy itself.

## IV. CONCLUSIONS

By characterising the topology of single-cell proliferation lineages, we showed that the heterogeneity observed in human BMSC clonal colonies is not due to random cell-to-cell variations, but rather to a correlated structure, which can only be due to inheritance. Our study therefore indicates that epigenetic hereditary factors play an important role in determining activity and cell-cycle exit in the early developmental stages of stem cell colonies.

The evidence of inheritance that we obtain from the inactivity imbalance is due to the existence of *intra-colony* heterogeneities: it is only thanks to the emergence throughout the lineage of changes in the propensity to become inactive that we can detect the anomalous imbalances that contribute to lowering the entropy; on the contrary, colonies with no inactive cells do not give any signal; as we have already noted, inheritance is undetectable in absence of mutations. Once the presence of inheritance is established through the intra-colony fluctuations, it is legitimate to infer that the same hereditary mechanism is responsible for the colony-level permanence of epigenetic traits, thus giving rise to *inter-colony* heterogeneities. Hence, our result supports the hypothesis that the strong colony-level heterogeneities met when dealing with in vivo transplant of BMSC cells — heterogeneities that strongly affects the effectiveness transplants and thus of skeletal regeneration therapies — do have an epigenetic hereditary origin.

The distribution of inactive cells across a colony determines the topology of the lineage and its relative inheritance entropy. Active cells, on the other hand, constitute a separate population that enters the entropy-based hereditary test only in a relatively passive way. Experimental evidence, though, shows that the topology of BMSC lineages (determined by inactive cells) is strongly correlated to the kinetics of the colony, i.e. to the statistics of the division times of the active cells [35]; more specifically, data show that colonies with the larger populations of inactive cells are also characterized by the slower populations of active cells [35]. This result suggests that not only cell-cycle exit, but also cell-cycle duration might be regulated by epigenetic hereditary mechanisms. To confirm this point, though, it is necessary to carefully study the correlation between the division times. This has been done extensively for bacterial colonies in relation to genetic factors affecting cycle time [33, 47], and to a scarcer degree for epigenetic factors in eukaryotic cells [7, 48], but never in the context of stem cells. Besides, in all past studies, the assessed correlation was not quite deep enough to trace in a robust way the hereditary ramifications of the division times across the lineage tree. This remains therefore an interesting open experimental and theoretical problem along the path to establishing epigenetic factors as the primary cause of the strong heterogeneity affecting stem cell colonies.

## V. ACKNOWLEDGEMENTS

We thank William Bialek, Fabio Cecconi and Edo Kussel for very insightful advice on the manuscript. ACa and TSG thank Enzo Branchini for discussions within the CC. ACa acknowledges the support and advice of the late Giovanni Cavagna. This work was supported by ERC Grant RG.BIO (Contract n. 785932), MIUR Grant INFO.BIO (Protocol n. R18JNYYMEY), and MIUR Grant PRIN2020 (Contract n. 2020PFCXPE-005).

## APPENDIX A: DETAILS ON THE LINEAGE SCRAMBLING PROCEDURE

The random scrambling of a biological lineage is a key step in the inheritance test, hence it has to be performed very carefully. The first important point is that the scrambled tree must have all potentially hereditary mother-daughter links severed, but at the same time it must belong to the same topological class as the original tree, namely it must have the same number of inactive cells *per generation* as the biological lineage; if this second requirement is violated we do not obtain a topologically homogeneous ensemble of trees, hence the entropy can drift to any value and no random-vs-biological comparison can be performed. To achieve these two conditions we proceed as follows (see Fig. 7).

We start the reshuffling at generation *k* = 2, since swapping the only two cells of generation *k* = 1 leaves the tree topologically unchanged. Two cells are randomly chosen among the 4 at *k* = 2 (red labels in Fig. 7) and these two cells are randomly swapped; when this is done, the descendants of the two cells are also swapped. Then we move to generation *k* = 3 and we do the same, i.e. we swap a pair of randomly chosen cells, exchanging also their sub-branches. This procedure is repeated recursively at all generations, each step of the procedure typically increasing the entropy, as any trace of non-random inheritance is deleted. It is crucial to swap only cells at the *same* generation, in order to keep the tree within the same topological class as the original biological one.

Even though the procedure that we have just described, and that is illustrated in Fig. 7, is the most intuitive one, exchanging just *one* pair of cells per generation makes the randomisation of the tree quite slow. To proceed more effectively in severing all potentially heritable connections in the tree, instead of just swapping two cells per generation, we perform a whole random *permutation* of all cells at each generation. As in the simple pair swap, also in the case of a permutation, whenever and wherever a cell is moved, it brings along with it all its descendants. The scrambling procedure of one tree is complete once a random permutation of all cells of each generation is performed.

## APPENDIX B: TOPOLOGICAL DISTANCE VS LINEAGE DISTANCE

Is it possible that the clustering of G_0_ cells on the same branches of the lineage tree, which we interpret as an hereditary character, is in fact an epiphenomenon of spatial correlation in the physical space of the dish? Namely, is it possible that chemicals/nutrient heterogeneities, local crowding, or other spatial factors might influence the propensity to enter the G_0_ phase in a way that is then reflected in the lineage topology? This is an old hypothesis, dating back to the sixties [49], shortly after the first evidence of correlations between division times of sister cells in bacterial colonies emerged [33], correlation that suggested to some authors that generation times could be affected by inherited factors [50].

Our data, however, show that our results are *not* due to spatial proximity, as we are going to explain below. In what follows we denote as “topological distance” the distance on the tree, namely the number of links separating two different cells in the lineage; on the other hand, the “physical distance” is the actual distance between the two cells on the dish.

Proximity in the physical space is not the same as proximity on the tree. More precisely, points that are close on the tree tend also to be close in physical space, since cells derived from a common close progenitor start their life closer in space than cells derived by far progenitors; but the crucial point is that the *vice-versa* is not true, namely points close in physical space are not necessarily close in the lineage (Fig. 8). For the sake of argument, let us assume that there exist on the Petri dish a localised spot of linear size *R* where spatial factors promote the G_0_ state; if a mitosis occurs in that G_0_-promoting spot and if the colony is already quite crowded — and hence mobility is limited — it is likely that also the two daughters’ mitosis will occur in that same spot. If these were the only mitosis taking place in the spot, this could explain the clustering that we observe in the lineage trees without the need to invoke an hereditary mechanism. However, this is not what happens, as clearly illustrated in Fig. 8: the range of topological distances between the points belonging to a physical region of size *R* is *very* wide; even a physical spot of a few hundred pixels, which is about the size of a cell, has a typical size on the tree of over 7 links, and it is very likely to find within this spot points with topological distance of up to 12 links, which is the maximum amplitude of any lineage. This result indicates that any hypothetical G_0_-promoting spatial spot — even a *very* small one — would contain many mitosis with *large* topological distances from each other, which would imply that the G_0_ cells emerging within the spot would be scattered all over the tree and not clustered within some branches, which is instead what we observe. We conclude that the G_0_ structure we discovered is very unlikely to be of spatial origin and that the hereditary explanation is the most plausible one.

**FIG. 8.**
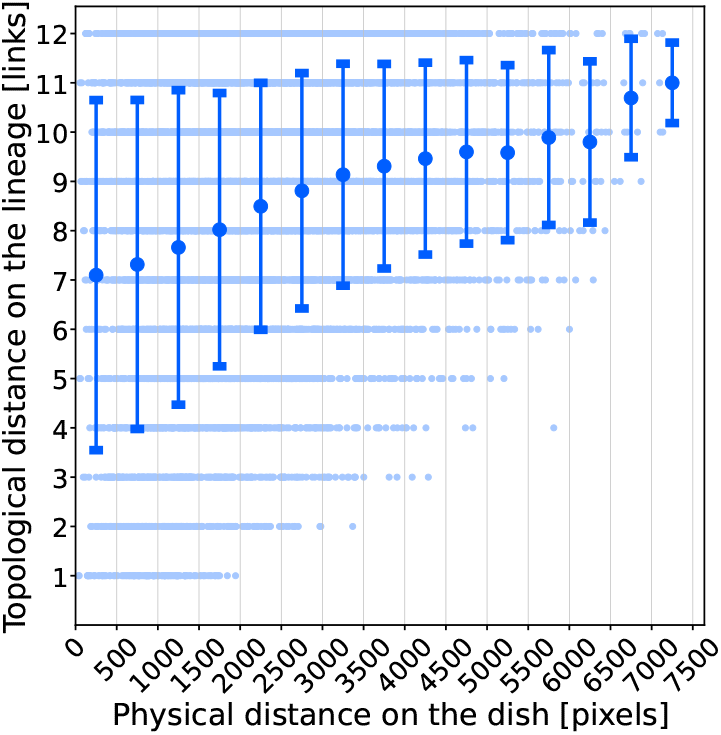
Mutual distances between all mitosis in a sample colony, `20241017_02` (light blue points); topological distance on the tree is measured in number of links separating two mitosis/nodes, while physical distance on the dish is measured in pixels separating the two mitosis. Correlation between the two types of distance is weak, with Spearman correlation coefficient *ρ* = 0.24 (P-value < 10^−6^). Even small physical distances correspond to a very wide range of topological distances; this is clear if we group together all physical distances in bins of 500 pixels and average in each bin the relative topological distances (dark large points — error bars are standard deviations). This plot demonstrates that short topological distances correspond to short physical distances, but that short physical distances do not correspond to short topological distances. In other words, a small spot on the dish is mapped onto a very large “region” on the tree: a spot of 250 pixels maps to an average distance on the tree of 7 links, which is very large in genealogical terms.

## Footnotes

1 This is a rather standard (and venerable) convention in the literature on cellular senescence: any difference between dead cells and cells that have ceased to divide is immaterial when dealing with population growth, see for example [36–38].

## References

[1] U. Deichmann, Developmental biology 416, 249 (2016).

[2] E. Jablonka and M. J. Lamb, Annals of the New York Academy of Sciences 981, 82 (2002).

[3] V. V. Lunyak and M. G. Rosenfeld, Human molecular genetics 17, R28 (2008).

[4] A. Meissner, Nature biotechnology 28, 1079 (2010).

[5] R. K. Ng and J. B. Gurdon, Cell cycle 7, 1173 (2008).

[6] E. H. Zion, C. Chandrasekhara, and X. Chen, Current opinion in cell biology 67, 27 (2020).

[7] O. Sandler, S. P. Mizrahi, N. Weiss, O. Agam, I. Simon, and N. Q. Balaban, Nature 519, 468 (2015).

[8] N. Mosheiff, B. M. Martins, S. Pearl-Mizrahi, A. Grünberger, S. Helfrich, I. Mihalcescu, D. Kohlheyer, J. C. Locke, L. Glass, and N. Q. Balaban, Physical Review X 8, 021035 (2018).

[9] S. Hormoz, Z. S. Singer, J. M. Linton, Y. E. Antebi, B. I. Shraiman, and M. B. Elowitz, Cell systems 3, 419 (2016).

[10] H. H. Chang, M. Hemberg, M. Barahona, D. E. Ingber, and S. Huang, Nature 453, 544 (2008).

[11] M. Plambeck, A. Kazeroonian, D. Loeffler, L. Kretschmer, C. Salinno, T. Schroeder, D. H. Busch, M. Flossdorf, and V. R. Buchholz, Proceedings of the National Academy of Sciences 119, e2116260119 (2022).

[12] M. Yampolskaya, L. Ikonomou, and P. Mehta, arXiv preprint 2506.04219 (2025).

[13] D. E. Wagner and A. M. Klein, Nature Reviews Genetics 21, 410 (2020).

[14] C. S. Baron and A. van Oudenaarden, Nature reviews molecular cell biology 20, 753 (2019).

[15] C. Weinreb, A. Rodriguez-Fraticelli, F. D. Camargo, and A. M. Klein, Science 367, eaaw3381 (2020).

[16] B. Raj, in Lineage Tracing: Methods and Protocols (Springer, 2025) pp. 299–310.

[17] B. Spanjaard, B. Hu, N. Mitic, P. Olivares-Chauvet, S. Janjuha, N. Ninov, and J. P. Junker, Nature biotechnology 36, 469 (2018).

[18] R. J. Ihry, K. A. Worringer, M. R. Salick, E. Frias, D. Ho, K. Theriault, S. Kommineni, J. Chen, M. Sondey, C. Ye, et al., Nature medicine 24, 939 (2018).

[19] M. Ogrodnik, Aging cell 20, e13338 (2021).

[20] C. M. Koch, S. Joussen, A. Schellenberg, Q. Lin, M. Zenke, and W. Wagner, Aging cell 11, 366 (2012).

[21] J. Franzen, A. Zirkel, J. Blake, B. Rath, V. Benes, A. Papantonis, and W. Wagner, Aging cell 16, 183 (2017).

[22] J. Franzen, T. Georgomanolis, A. Selich, C.-C. Kuo, R. Stöger, L. Brant, M. S. Mulabdić, E. Fernandez-Rebollo, C. Grezella, A. Ostrowska, et al., Communications biology 4, 598 (2021).

[23] D. Nachtwey and I. Cameron, in Methods in Cell Biology, Vol. 3 (Elsevier, 1969) pp. 213–259.

[24] T. H. Cheung and T. A. Rando, Nature reviews Molecular cell biology 14, 329 (2013).

[25] C. Behl and C. Ziegler, in Cell aging: Molecular mechanisms and implications for disease (Springer, 2013) pp. 9–19.

[26] M. M. Mens and M. Ghanbari, Stem cell reviews and reports 14, 309 (2018).

[27] S. Mareddy, R. Crawford, G. Brooke, and Y. Xiao, Tissue engineering 13, 819 (2007).

[28] S. Mareddy, N. Dhaliwal, R. Crawford, and Y. Xiao, Tissue Engineering Part A 16, 749 (2010).

[29] P. Bianco and P. G. Robey, Development 142, 1023 (2015).

[30] P. Bianco, X. Cao, P. S. Frenette, J. J. Mao, P. G. Robey, P. J. Simmons, and C.-Y. Wang, Nature medicine 19, 35 (2013).

[31] T. Kuczek and D. E. Axelrod, Mathematical biosciences 79, 87 (1986).

[32] D. A. Rennerfeldt and K. J. Van Vliet, Stem cells 34, 1135 (2016).

[33] E. Powell, Biometrika 42, 16 (1955).

[34] A. Genthon, T. Nozoe, L. Peliti, and D. Lacoste, PRX Life 1, 013014 (2023).

[35] A. Allegrezza, R. Beschi, D. Caudo, A. Cavagna, A. Corsi, A. Culla, S. Donsante, G. Giannicola, I. Giardina, G. Gosti, et al., arXiv preprint arXiv:2504.21818v2 (2025).

[36] T. Kirkwood and R. Holliday, Journal of Theoretical Biology 53, 481 (1975).

[37] R. Holliday, L. Huschtscha, G. Tarrant, and T. Kirkwood, Science 198, 366 (1977).

[38] C. B. Harley and S. Goldstein, Science 207, 191 (1980).

[39] S. E. Luria and M. Delbrück, Genetics 28, 491 (1943).

[40] W. A. Rosche and P. L. Foster, Methods 20, 4 (2000).

[41] D. H. Colless, “Phylogenetics: the theory and practice of phylogenetic systematics,” (1982).

[42] M. Fischer, L. Herbst, S. Kersting, A. L. Kühn, and K. Wicke, Tree balance indices: A comprehensive survey (Springer Nature, 2023).

[43] S. F. Gull and G. J. Daniell, Nature 272, 686 (1978).

[44] H. Theil, Economics and information theory (Amsterdam: North-Holland,, 1967).

[45] M. K. Strasser, P. S. Hoppe, D. Loeffler, K. D. Kokkaliaris, T. Schroeder, F. J. Theis, and C. Marr, Nature communications 9, 2697 (2018).

[46] W. Wagner, P. Horn, M. Castoldi, A. Diehlmann, S. Bork, R. Saffrich, V. Benes, J. Blake, S. Pfister, V. Eckstein, et al., PloS one 3, e2213 (2008).

[47] R. Staudte, M. Guiguet, and M. C. d’Hooghe, Journal of theoretical biology 109, 127 (1984).

[48] S. Hormoz, N. Desprat, and B. I. Shraiman, Proceedings of the National Academy of Sciences 112, E2281 (2015).

[49] G. Froese, Experimental cell research 35, 415 (1964).

[50] H. Kubitschek, Experimental cell research 26, 439 (1962).

